# Traction Force and Mechanosensing can be Functionally Distinguished Through the Use of Specific Domains of the Calpain Small Subunit

**DOI:** 10.1101/2023.03.07.531592

**Authors:** Bingqing Hao, Karen A. Beningo

## Abstract

Cell migration is a fundamental process pertaining to many critical physiological events. The ability to form and release adhesion structures is necessary for cell migration. The Calpain family of cysteine proteases are known to target adhesion proteins as their substrates and modulate adhesion dynamics. The two best studied Calpains, Calpain 1 and Calpain 2 form catalytically active holoenzymes through heterodimerization with a common non-catalytic regulatory small subunit known as Calpain 4. In previous studies, we determined that calpains are important in the production of traction forces and in the sensing of localized mechanical stimulation from the external environment. We found that perturbation of either Calpain 1 or 2 had no effect on the generation of traction forces. However, traction forces were weak when Calpain 4 was silenced. On the other hand, silencing of Calpain 1, 2, or 4 resulted in deficient sensing of external mechanical stimuli. These results together suggest that Calpain 4 functions independent of the catalytic large subunits in the generation of traction forces but functions together with either catalytic subunit in sensing external mechanical stimuli. The small subunit Calpain 4 contains 268 a.a. and is composed of 2 domains, the N-terminal domain V and C-terminal domain VI. Domain VI is a calmodulinlike domain containing five consecutive EF-hand motifs, of which the fifth one heterodimerizes with a large subunit. Moreover, domain V contains the common sequence GTAMRILGGVI that suggests cell membrane interactions. Given these attributes of domain V and VI of Calpain 4, we speculated that an individual domain might provide the functional properties for either traction or sensing. Therefore, each domain was cloned and expressed individually in *Capn4−/−* cells and assayed for traction and sensing. Results revealed that over-expression of domain V was sufficient to rescue the traction forces defect in *Capn4−/−* cells while overexpression of domain VI did not rescue the traction force. Consistent with our hypothesis, overexpression of domain VI rescued the sensing defect in *Capn4−/−* cells while overexpression of domain V had no effect. These results suggest that individual domains of Calpain 4 do indeed function independently to regulate either traction force or the sensing of external stimuli. We speculate that membrane association of Calpain 4 is required for the regulation of traction force and its association with a catalytic subunit is necessary for mechanosensing.

## Introduction

Cell migration has been implicated in many critical biological processes, including embryonic development, wound healing, immunological responses, and cancer metastasis. A coordinated series of events are required for cell migration, including: protrusion at the cells front edge, adhesion of the protruded area to the substrate, pulling of the cell body, and retraction at the cell rear (Friedl and Alexander, 2011; Ridley, 2003). These processes require the functions of focal adhesions to transmit the traction forces exerted onto the substrate by cells and to sense the mechanical signals from the external environment (Flevaris, et al., 2007; Hynes, 2002; Lauffenburger and Horwitz, 1996; Ridley, 2003).

Focal adhesions are dynamic assemblies of adaptor proteins and integrin transmembrane receptors that couple the extracellular matrix (ECM) to the actin cytoskeleton. Members of the calpain family of calcium dependent cysteine proteases have long been implicated in the turnover of focal adhesion component proteins (Bhatt et al., 2002; Dourdin et al., 2001; Franco et al., 2004b; Goll et al., 2003). The two isoforms μ-calpain and m-calpain are the most well characterized members of this family. These two catalytic domains share a common 28 kDa small subunit, known as calpain 4 (CAPNS1 OR CAPN4, encoded by *Capn4* gene), which heterodimerizes with distinct 80 kDa large subunits known as calpain 1 and calpain 2 (CAPN1 AND CAPN2, encoded by *Capn1 and Capn2* genes respectively). Structurally, the protease domains are only located within the large subunits but are absent in the small subunit. There are two terminal domains that make up the small subunit, also known as the regulatory subunit: the NH_2_-terminal domain V, and the COOH-terminal domain VI (Goll et al., 2003). Domain V is Gly rich and contains a potential phospholipid binding region GTAMRILGGVI (Crawford, 1990; Daman, 2001; Imajoh et al., 1986). Domain VI contains five EF-hand motifs with the fifth EF hand interacting with the corresponding fifth EF hand from domain IV of the large subunit for assembly of the holoenzyme (Franco and Huttenlocher, 2005).

The role of calpains in cell migration has been widely investigated. Inhibition of calpain results in reduced cell migration, delayed retraction of the cell’s rear, inhibition of focal adhesion disassembly and translocation, impairment of cell spreading, and modulation of cancer cell invasion (Bhatt, 2002; Huttenlocher, 1997; Mamoune, 2003; Potter, 1998). In a study seeking the functions for calpain isoforms in regulating membrane protrusion, it was found that despite the absence of obvious defects in *Capn1* silenced cells, silencing of *Capn2* resulted in protrusion defects and delayed focal adhesion disassembly (Franco et al., 2004a; Franco et al., 2004b).

Although much attention has been given to the functions of calpains on cell migration, the calpain small subunit has been under investigated as it has been mainly associated with a regulatory role in the holoenzymes (Goll et al., 2003). *Capn4^−/−^* embryonic fibroblasts display a reduced rate of cell migration, abnormal organization of focal adhesions with a loss of centralized focal adhesions, and delayed retraction of membrane projections, which suggests a deficiency in focal adhesion maturation and turnover (Dourdin, 2001). We explored the function of calpains in the generation of traction forces and mechanosensing, and discovered that traction force was inhibited by the disruption of *Capn4* expression, but not by the inhibition of the large subunits or the overexpression of calpastatin (an endogenous inhibitor of the catalytic subunits). On the other hand, inhibiting either large subunit or interrupting the small subunit led to defects in the mechansensing to the localized force and substrate topography. Consistently *Capn4^−/−^* cells have abnormal stress fibers and a reduced number of stress-fiber-associated, vinculin-containing adhesions (Undyala, 2008). These results implicate only the calpain small subunit in the regulation of traction forces but both large and small subunits in mechanosensing.

In this study, we performed a simple domain function analysis with the hypothesis that calpain 4 would have one domain that coordinates mechanosening and a second domain involved in traction force. We overexpressed either domain V or domain VI of calpain 4 in *Capn4^−/−^* cells and looked for phenotypic changes in migration, adhesions, traction force and mechanosensing. We discovered that overexpression of domain V rescued the defective production of traction force and abnormal focal adhesion organization in *Capn4^−/−^* cells and promotes cell migration rates. On the other hand, overexpression of domain VI restored both the ability to sense the localized mechanical force and the protease activity in *Capn4^−/−^* cells. These results suggest that the calpain small subunit has a protease-independent function in the production of traction force through domain V, and a mechanosensing function specific to domain VI. To our knowledge this is the first example of a single molecule possessing the ability to regulate traction with one protein domain and mechanosensing with a second independent domain without interfering with the activities of the other domain.

## EXPERIMENTAL PROCEDURES

### Cell Culture

Mouse embryonic fibroblasts expressing a defective small calpain subunit have been described previously (Arthur et al., 2000; Dourdin et al., 2001), and are referred to as *Capn4^−/−^* cells in this study. Mouse embryonic fiborolasts (MEF) and *Capn4^−/−^* cells were used in this study. MEFs were purchased from ATCC. Cells were maintained in Dulbecco’s Modified Eagle’s Medium-high glucose (Sigma) supplemented with 10% fetal bovine serum (FBS) (Hyclone) and 1% penicillin/streptomycin/glutamine (Gibco) and incubated at 37 °C under 5% CO2 in a humidified cell culture incubator. Cells were passed by trypsinization using 0.1% trypsin-EDTA (0.25% trypsin-EDTA diluted with HBSS, Gibco). Trypsinization was stopped by adding complete media. The passage number of either cell type never exceeded six passages.

### Cloning of Domain V and VI of Capn4 and DNA Constructs

The pAcGFP1-N1 (Clontech) was transformed into E.coli and purified with an E.Z.N.A Plasmid Mini Kit (Omega). Sequences of domain V, VI or full length *Capn4* were amplified by PCR from a pEGFP-capn4 under the following conditions: 30 cycles of 98 °C for 10 sec followed by 68 °C for 1 min using PrimeSTAR HS DNA Polymerase with GC buffer (Takara, R044A). The primers used were as follows: full length *capn4* was amplified with the forward primer 5’-ACCGCTCGAGATGTTCTTGGTGAATTCGTT-3’ and the reverse primer 5’-ATCGGGATCCGCGGAATACATAGTCAGCTGCA-3’; domain V was amplified with the forward primer 5’-ACCGCTCGAGATGTTCTTGGTGAATTCGTT-3’ and the reverse primer 5’-TACGGGATCCGCGAACTGACGGACTTCTTCA-3’; and domain VI was amplified with the forward primer 5’-ACCGCTCGAGATGAGGAAACTTTTTGTCCAG-3’ and the reverse primer 5’-ATCGGGATCCGCGGAATACATAGTCAGCTGCA-3’. PCR products were resolved on 1% agarose gels and visualized by ethidium bromide (1% solution, Fisher) staining. The resolved bands were then purified using a Qiaquick gel extraction kit (Qiagen, 28706). Purified PCR products and pAcGFP1-N1 were incubated with XhoI and BamHI (New England Biolabs) under 37 °C for 4 hrs in 1X buffer 3 supplemented with 1% BSA. The double digested PCR products and plasmid were again purified with the Qiaquick gel extraction kit. To insert either domain V or VI into pAcGFP1-N1, ligation of double digested fragment of either domain with double digested pAcGFP1-N1 was performed with the LigaFAst Rapid DNA Ligation System (Promega, M8226) following the manufacturer’s suggested protocol. These constructs were transformed into E. coli to collect plasmids, and successful insertions were confirmed by sequencing (Applied Genomics Technology Center, Wayne State University).

### Nucleofection of Capn4^−/−^ Cells and Overexpression of Domains

Nucleofection was performed using the Amaxa MEF2 Nucleofector Kit (Lonza) following the manufecturer’s suggested protocol. Briefly, *Capn4^−/−^* cells were trypsinized with 0.1% Trypsin-EDTA and collected by centrifuging at 2000 rpm for 5 min. Collected cells were then resuspended in an appropriate volume of the mixture of the included MEF 2 nucleofector solution and supplement 1 followed by adding up to 5 ug of the prepared plasmid. The total volume of the MEF 2 Nucleofector solution and supplement 1 mixture and the plasmid added up to 100 μl, which was mixed well and transferred to an electroporation cuvette. The cuvette was then inserted into the Nucleofector II system (Amaxa) and the program MEF A-023 was run. 500μl of RPMI-1640 medium (Sigma) was immediately added. Nucleofected cells were then seeded according to the requirement of the following procedures.

### Protein Extraction and Western Blotting

Proteins were extracted from each cell line with triple detergent lysis buffer (TDLB): pH8 50 mM Tris HCl, 150 mM NaCl, 1% NP-40, 0.5% sodium deoxycholate, 0.1% SDS, into which Protease Inhibitor Cocktail (Sigma) and HaltTM Phosphatase Inhibitor Cocktail were added (Thermo). An 80% confluent 100 mm culture dish (Nunc^TM^) was placed on ice and washed twice with ice-cold phosphate-buffered saline (PBS) followed by 25 min of incubation with 300 μl TDLB on ice. Lysed cells were collected by an ice-cold cell lifter and centrifuged at 13,000 rpm for 10 min to get rid of cell debris. Protein concentration was measured by Lowry high method with the Bio-Rad DC protein assay kit. 20 μg of proteins from each cell line were loaded into a 4-20% gradient Tris– HEPES–SDS precast polyacrylamide gel system (Pierce) and resolved at 100 V for 1 hour. Proteins were then transferred onto an Immun-Blot PVDF Membrane (Bio-Rad) using a Trans-blot SD Semi-dry Transfer Cell (Bio-Rad) at 20 V for 30 min. Following transfer, the membrane was blocked for 1 hour using 5% milk in Tris Buffered Saline – 0.1% Tween (0.1% TBS/T) and then probed with the primary antibody. Primary antibody for GFP (sc-8334, Santa Cruz) was diluted at 1:500 in 5% milk in 0.1% TBS/T and incubated at 4 °C overnight with mild agitation. After washing 20 min for 3 times with 0.1% TBS/T, the secondary antibody HRP-linked Rabbit IgG (NA934, Amersham) was diluted at 1:10,000 in 5% milk in 0.1% TBS/T and incubated for 1 hour at room temperature. After washing 20 min for 3 times, the membrane was detected using ECL Plus Western Blotting Detection Reagents (Amersham).

### Preparation of Polyacrylamide Substrates

A series of polyacrylamide substrates of different stiffnesses were prepared as described previously (Beningo et al., 2002). Briefly, a flexible 75 μm x 22 mm polyacrylamide substrate was made in a cell culture chamber dish in which 0.2 μm fluorescent microbeads were embedded. The acrylamide (acryl, Bio-rad) concentration was fixed at 5% while N,N-methylene-bis-acrylamide (bis, Bio-rad) varied from 0.04% to 0.1% to attain different stiffnesses of the substrates. Traction force microscopy (TFM) was performed with the 5%/0.08% Acryl/Bis substrates and the mechanosensing assay to applied forces was performed with 5%/0.1% Acryl/Bis substrates. The substrates were then coated with 5 μg/cm^2^ fibronectin (Sigma) at 4^°^C overnight. Cells were seeded onto the substrates overnight prior to TFM or mechanosensing.

### Traction Force Microscopy

After an overnight incubation in the cell culture incubator, images for cells were collected as described previously (Beningo et al., 2002). Briefly, 3 images were taken for a single cell under 40X objective lens: a bright field image of the cell, an image for the fluorescent beads with the cell on the substrate, and another image for the fluorescent beads after the cell was removed by a pointed microneedle. Bead displacement with or without the cell and the cell and nuclear boundaries calculated by DIM (Yu-li, Wang) were used to generate and render traction stress values by using a custom made algorithm provided by Dr. Micah Dembo (Boston University) as described previously (Dembo and Wang, 1999; Marganski et al., 2003). Images of 12-18 cells for each cell line were collected.

### Mechanosensing Assay to Applied Mechanical Stimulation

Flexible polyacrylamide substrates of 5/0.1% Acryl/Bis coated with fibronectin were prepared as described above. Cells were seeded onto the substrates and allowed to adhere overnight under regular cell culture conditions. As described previously (Lo et al., 2000), a cell was monitored for 10 min for its migration trajectory before a blunted microneedle was pressed onto the substrate in front of the direction the cell was migrating to generate a pushing force onto the cell. The pushing force will release the tension on the substrate. Images were taken every 3 min for 1 hour. If a cell responds to the pushing force by avoiding it, a “1” is recorded; if a cell continues to migrate on the same trajectory, a “0” is recorded. For each cell line, 12-18 cells were observed.

### Immunofluorescence

After being flamed, No. 1.5 glass coverslips (Fisher) were coated with 5 μg/cm^2^ fibronectin (Sigma) at 4°C overnight and blocked with 1% BSA in PBS at 4°C overnight. Cells were seeded onto the glass coverlips and allowed to attach overnight under regular cell culture conditions. The cells were then fixed and permeabilized with the following steps: first incubate for 10 min with 2.5% paraformaldehyde in 1X PBS at 37^°^C; then incubate with 2.5% paraformaldehyde in 1X PBS with 0.1% Triton X-100 at 37^°^C; followed by incubation of 5 min with 0.5 mg/ml NaBH4 solution. After fixation and permeabilization, cells were blocked with 5% BSA in PBS for 1 hour at room temperature, and then incubated with anti-vinculin antibody (Sigma, V4505) at a 1:200 dilution for 3 hours at room temperature. Following 3 washes of 15 min, Alexa Fluor^®^ 546 anti-mouse secondary antibody was added at a 1:500 dilution in 5% BSA for an incubation of 1 hour at room temperature. After the final washes (3 x 15 min each), mounting media (ph7.8, 0.1% PPD, 1X PBS, 50% glycerol, 30% Q-H_2_O) was added. Images were taken with appropriate filters for both GFP and RFP signals. The number and size of vinculin containing plaques were measured using the NIH Image J (NIH).

### Calpain Activity Assay

Calpain activity was quantified using a calpain activity fluorometric assay kit (Biovision) following the manufacturer’s instructions, except using a modified lysis buffer. Briefly, cells were lysed with TDLB as described above, into which Protease Inhibitor Cocktail (Sigma) and HaltTM Phosphatase Inhibitor Cocktail were added (Thermo). The protein concentration was calculated by Lowry high method with the Bio-Rad DC protein assay kit. 50 μg of cell extracts was mixed and incubated with the reaction buffer and calpain substrate Ac-LLY-AFC provided by the kit for 1 hour at 37°C in the dark. The samples were then transferred to a 96-well plate, and the reactions were measured at 400/505 nm with a Spectramax Gemini Fluorescence Luminescence Microplate Reader (Molecular Devices, Sunnyvale, CA).

### Cell Migration Assay

Glass coverslips were coated with 5μg/cm^2^ fibronectin (Sigma) at 4°C overnight, then cells were seeded and allowed to attach overnight under regular cell culture conditions. The migration trajectory of a single cell was observed for 2 hours at 2 min intervals with a 40X objective lens. All the images were analyzed with the custom built dynamic image analysis system software (DIM, Y-L. Wang) to calculate the linear speed and persistence of each cell. for each cell line, 10-15 cells were observed.

### Microscopy

Images for all experiments described above were acquired with an Olympus IX81 ZDC inverted microscope fitted with a custom-built stage incubator to maintain cells at 37°C under 5% CO_2_ for live cell imaging and a SPOT Boost EM-CCD-BT2000 back-thinned camera (Diagnostic Instruments Inc., Sterling Heights, MI). The camera was driven by the IPLab software (BD Biosciences).

## RESULTS

### Plasmid Construction and Overexpression of Capn4 Domains in Capn4^−/−^ Cells

Calpain 4 regulates the generation of traction forces in MEF cells in addition to the canonical regulatory function for the holoenzyme (Undyala et al., 2008). Previous research showed that the generation of traction forces was inhibited by the disruption of *Capn4* expression but not by the knock-down of *Capn1* or *Capn2* or even the overexpression of calpastatin, the endogenous calpain inhibitor. However, localized tension sensing required the function of both large and small subunits of the holoenzyme (Undyala et al., 2008). To further evaluate the functions of domain V (DV) and VI (DVI) on the generation of traction forces and mechanosensing, each domain or the full-length *Capn4* was first cloned into the pAcGFP1-N1 and then overexpressed in *Capn4^−/−^* cells (S1, *A*). The overexpression of each plasmid was confirmed by Western blots (S1, *B*). Successful overexpression of the *Capn4* domains in *Capn4^−/−^* cells make it possible to test the impact of either *Capn4* domain on the cell’s ability to generate traction forces and sense the external stimulus.

### Overexpression of DV Rescues the Defect of Traction Force Generation in Capn4^−/−^ Cells

Previous studies on the function of the calpain small subunit in cell migration revealed reduced production of traction forces in Capn4^−/−^ cells compared to wild-type MEF cells, while inhibition of *Capn1* or *Capn2* or the overexpression of calpastatin did not affect the production of traction forces (Undyala et al., 2008). To understand the function of each domain of the calpain small subunit on traction force production in migrating fibroblasts, *Capn4^−/−^* cells expressing either the DV or DVI plasmid were plated on flexible polyacrylamide substrates covalently coated with fibronectin followed by traction force microscopy (TFM) (Dembo & Wang, 1999). Traction forces were calculated based on the magnitude of bead displacement within the substrate with or without the attached cell, and then vector maps were generated (Fig. 1 *A*). The magnitude of the traction forces produced in *Capn4^−/−^* cells expressing either DV, DVI, or full-length *Capn4* gene were compared with wild-type MEFs and *Capn4^−/−^* cells.

**FIGURE 1.**
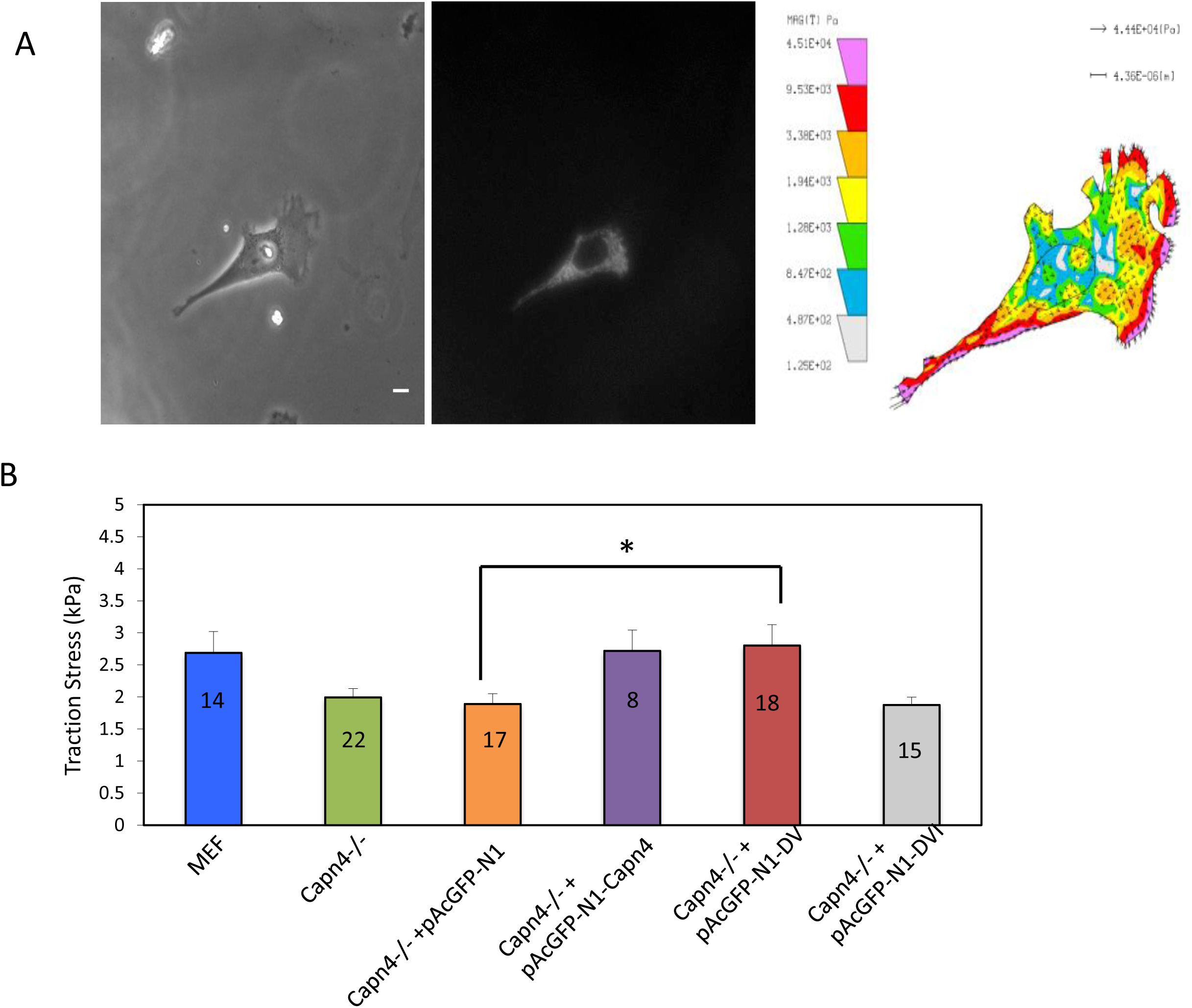
Overexpression of DV rescues the defect of traction force production in *Capn4^−/−^* cells. *A*, Cells were seeded onto flexible polyacrylamide substrates coated with fibronectin and allowed to attach overnight. Images of the embedded fluorescent microbeads with or without the cell were taken for each cell. The change in bead position (displacements) and the cell and nuclear boundary were used to calculate traction stress when incorporated into a custom made algorithm. The vector plot on the right indicates the magnitude and direction of traction stress exerted by a single cell, that on the left shows a cell. In these vector maps, arrowheads indicate direction and magnitude of forces. Red and pink highlight areas of strongest force and blue and gray indicate regions of weaker force as indicated on the color bar (Mag. bar = 10μm). B, The bar graph indicates the average traction stress (force/area) exerted by these cell lines: MEFs, *Capn4^−/−^* cells, *Capn4^−/−^* cells expressing the empty plasmid pAcGFP1-N1, *Capn4^−/−^* cells expressing full-length *Capn4* gene, *Capn4^−/−^* cells expressing DV, and *Capn4^−/−^* cells expressing DVI. The number of cells recorded for each cell line is marked on top of the respective bar. Statistical analysis was performed by student’s t-test. * indicates p<0.05; NS indicates a nonsignificant.

Compared to wild-type MEF cells (avg. 2.69kPa), *Capn4^−/−^* cells produced significantly reduced level of traction forces (avg. 1.99 kPa, *p*=0.03) (Fig. 1, *B*), which is consistent with the previous study (Undyala et al., 2008). Moreover, the expression of the full-length *Capn4* restored the ability to produce traction forces in *Capn4^−/−^* cells (avg. 2.72 kPa, *p*=0.0.03), and the expression of the empty plasmid had no effect (avg. 1.89 kPa). Meanwhile, expression of DV in *Capn4^−/−^* cells rescued the traction force production to a similar magnitude as in MEF cells (avg. 2.80 kPa, *p*=0.02), while *Capn4^−/−^* cells expressing DVI only produced traction forces at a similar magnitude to *Capn4^−/−^* cells (avg. 1.87 kPa). These results suggest that expression of DV in *Capn4^−/−^* cells is sufficient to rescue the traction force production defect, and that generation of these forces is mainly mediated through DV of calpain 4 but not DVI.

### The Deficient Mechanosensing in Capn4^−/−^ Cells is Rescued by Overexpressing DVI

Cells sense the mechanical signals from the extracellular environment including the substrate stiffness, topography, and localized mechanical stimuli, and couple these signals with mechanosensitive changes in the cytoskeletal networks, interaction with the extracellular matrix (ECM), and cellular force production (Engler et al., 2006; Guilak et al., 2009; Liedtke & Kim, 2005; Menon and Beningo, 2011). In previous research, our lab tested various calpain deficient cells for their ability to respond to localized mechanical stimuli in an assay where cells were seeded onto polyacrylamide substrates and a blunted needle was used to push on the substrate against the direction the cell was migrating. A wild-type MEF cell responds to the localized pushing force by avoiding it. However, *Capn4^−/−^* cells were deficient in sensing the applied force. *Capn1*, *Capn2*, or *Capn4* deficient were found to be unresponsive to the localized pushing force (Undyala et al., 2008).

Given the fact that DV of calpain 4 rescues the defects of traction force production in *Capn4^−/−^* cells, we asked whether any of the calpain 4 domains are also involved in the mechanosensing process. *Capn4^−/−^* cells expressing DV or DVI, wild-type MEFs and *Capn4^−/−^* cells were observed when the localized stimulus was applied, and data was recorded as either “1” for responding or “0” for non-responding. Data showed that expression of DVI restored the mechanosensing defect in *Capn4^−/−^* cells to the level of MEF cells, although expression of DV was not able to restore the mechanosensing ability in *Capn4^−/−^* cells. Unlike *Capn4^−/−^* cells expressing DVI, *Capn4^−/−^* cells and control *Capn4^−/−^* cells expressing an empty GFP plasmid were unable to sense the localized pushing force (Fig. 2, *A, B*). These results suggest that instead of DV, the function of sensing the localized pushing force is mediated through DVI of the calpain small subunit.

**FIGURE 2.**
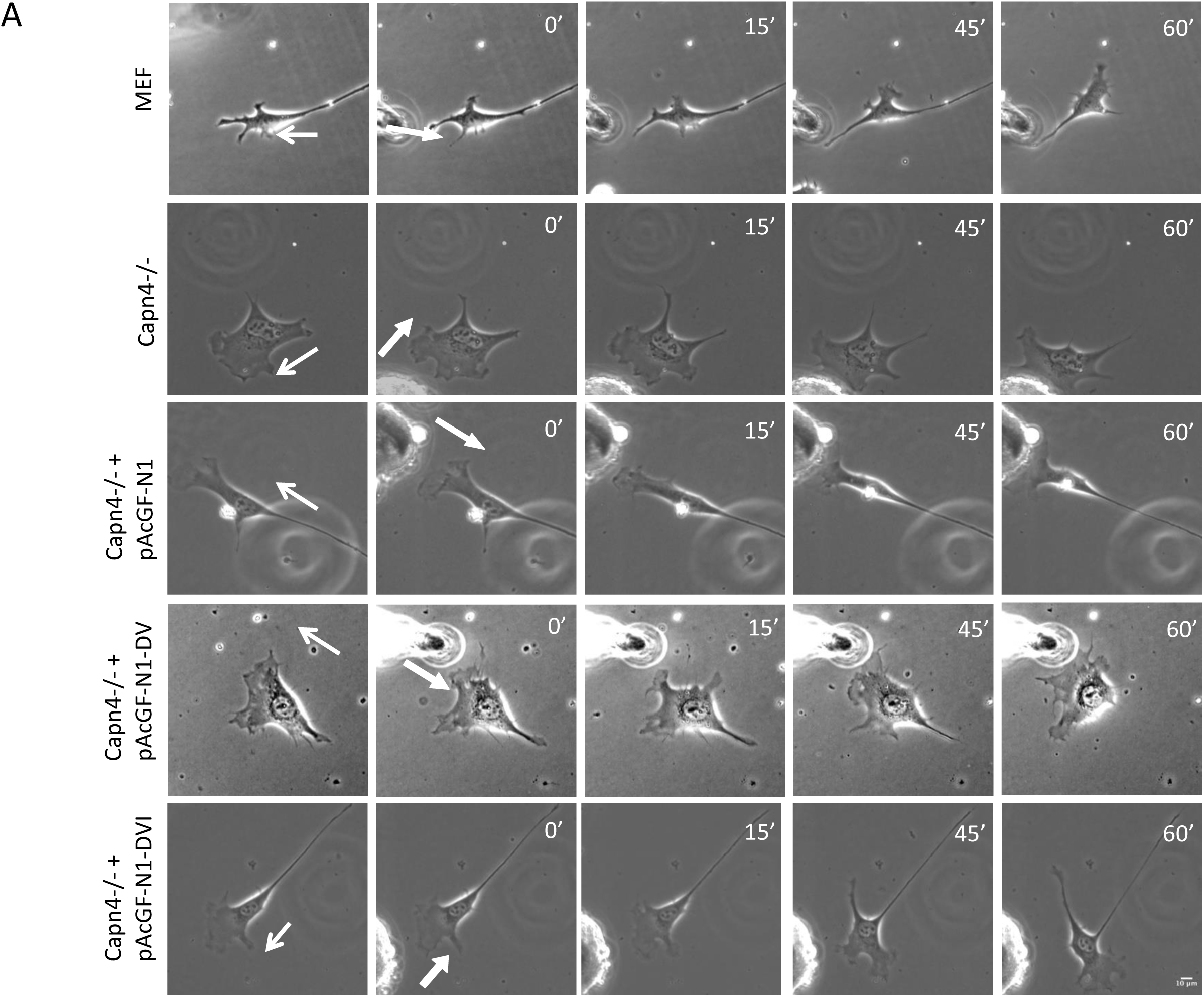
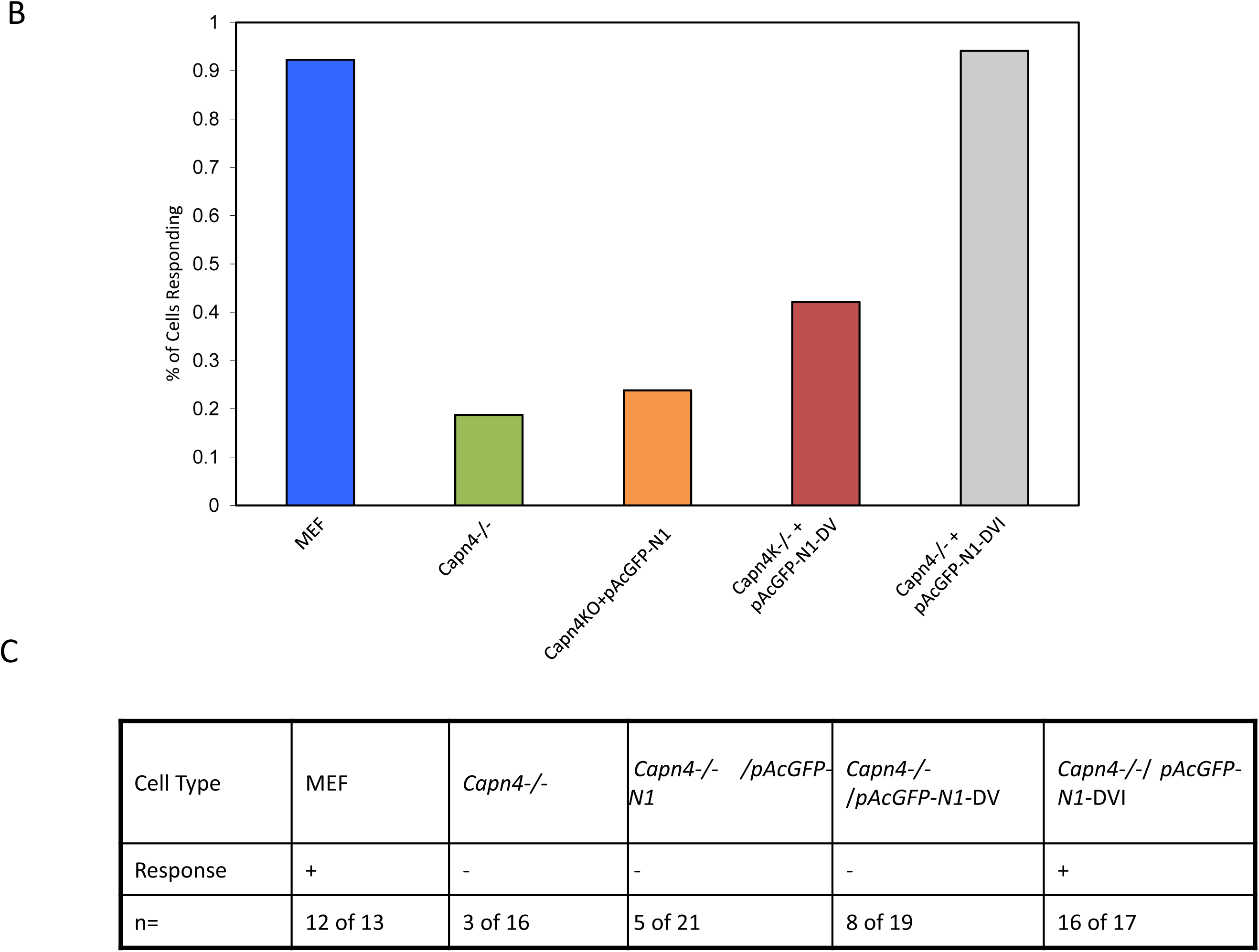
Overexpression of DVI in *Capn4^−/−^* cells restores the ability to sense the localized stimulus. *A*, Representative time-lapse images show the responses of cells to the applied localized stimulus. The included cells lines are: a MEF cell (top row), *Capn4^−/−^* cells (the second row); *Capn4^−/−^* cells expressing pAcGFP-N1 (the third row); *Capn4^−/−^* cells expressing DV (the fourth row), and *Capn4^−/−^* cells expressing DVI (the bottom row). The thin arrow denotes the original migration direction of the cell, and the thick arrow denotes the direction of the pushing force by the blunted needle (Mag. bar = 10μm). *B*. The bar graph indicates the percentage of cells responding to the localized stimulus. The observed cell lines are: MEFs, *Capn4^−/−^* cells, *Capn4^−/−^* cells expressing pAcGFP1-N1, *Capn4^−/−^* cells expressing full-length *Capn4*, *Capn4^−/−^* cells expressing DV, and *Capn4^−/−^* cells expressing DVI. *C*. The table summarizes the responses of cells for each cell line. “+” represents a positive reaction and “-” represents a negative reaction. The numbers of the representative cells for each cell line are also listed in this table. As expected, *Capn4^−/−^* cells displayed deficient mechanosensing compared to MEFs. In comparison to *Capn4^−/−^* cells expressing DV that are deficient in mechanosensing, *Capn4^−/−^* cells expressing DVI sensed the stimulus from the external environment as well as MEFs.

### Overexpression of DV Promotes the Maturation of Focal Adhesions

Traction forces are exerted onto the substrate through focal adhesions, which connect the actin cytoskeleton to the extracellular matrix (ECM) (Chrzanowska-Wodnicka and Burridge, 1996; Hotulainen and Lappalainen, 2006; Schoenwaelder and Burridge, 1999). *Capn4^−/−^* cells were previously found to have distinct morphology, including a loss of central focal adhesions, stabilization of focal complexes at the cell periphery, and fewer and less prominent actin stress fibers compared to wild-type MEFs. The same phenomena were not observed in *Capn1-* and *Capn2-* knockdown cells (Dourdin et al., 2001; Undyala et al., 2008).

Focal adhesions are dynamic structures. Nascent focal adhesions originate in lamellipodium. While the sizes of many focal adhesions continue to increase as they mature into the center of a cell and become larger plaques, others may simply disassemble (Alexandrova et al., 2008; Beningo et al., 2001; Mathew et al., 2011; Papusheva and Heisenberg, 2010). The Lack of centralized focal adhesions suggests a perturbation in the focal adhesion maturation process in *Capn4^−/−^* cells.

To determine whether expressing DV in *Capn4^−/−^* cells changes the focal adhesion organization, *Capn4^−/−^* cells expressing either DV or DVI, MEFs, and *Capn4^−/−^* cells were seeded onto fibronectin coated glass coverslips, fixed with paraformaldehyde and immuno-stained with anti-vinculin antibody and fluorescently labeled secondary antibodies. As expected, MEFs displayed normal focal adhesion organization localized to both the cell center and periphery in contrast to *Capn4^−/−^* cells where a loss of centralized focal adhesions with prominent vinculin containing focal adhesions located at the cell periphery. Furthermore, expression of DV in *Capn4^−/−^* cells rescued the abnormal organization of focal adhesions with many found in the center of cells, but this was not observed in *Capn4^−/−^* cells expressing DVI (Fig. 3, *A*). Also, quantification of the size and number of focal adhesions in each cell line displayed a significant decrease (*p*=0.0007) in the number of adhesions sizing from 0.5 to 1.5 sq.μm (nascent adhesions) in *Capn4^−/−^* cells. Meanwhile, expressing DV in *Capn4^−/−^* cells increased the number of nascent adhesions significantly (*p*=0.005), although not completely restored to the level of MEF cells (Fig. 3, *B*). However, *Capn4^−/−^* cells expressing DVI showed no significant increase in the number of nascent adhesions compared to *Capn4^−/−^* cells. For focal adhesions with a size smaller than 0.5 sq.μm, the only significant difference was between *Capn4^−/−^* cells and *Capn4^−/−^* cells expressing DV (Fig. 3, *B*). When measuring focal adhesions larger than 1.5 sq.μm, no significant difference is observed between any two cell lines, although *Capn4^−/−^* cells expressing DV do have elevated quantities of focal adhesions larger than 1.5 sq.μm. Altogether, these results suggest that in addition to restoring the production of traction forces in *Capn4^−/−^* cells, expression of DV, but not DVI, rescues the abnormal focal adhesion organization defects observed in *Capn4^−/−^* cells, and also aid in their maturation.

**FIGURE 3:**
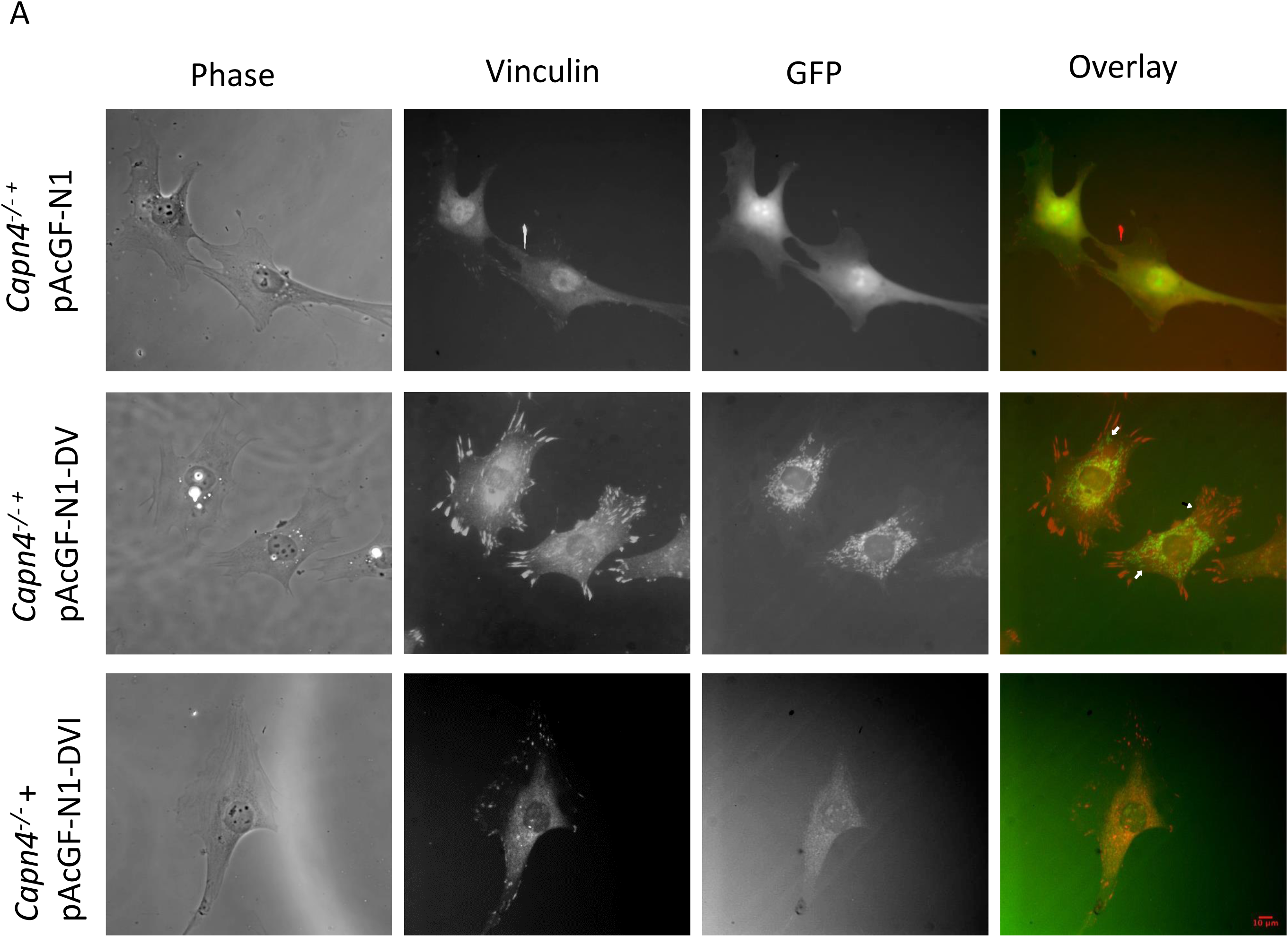
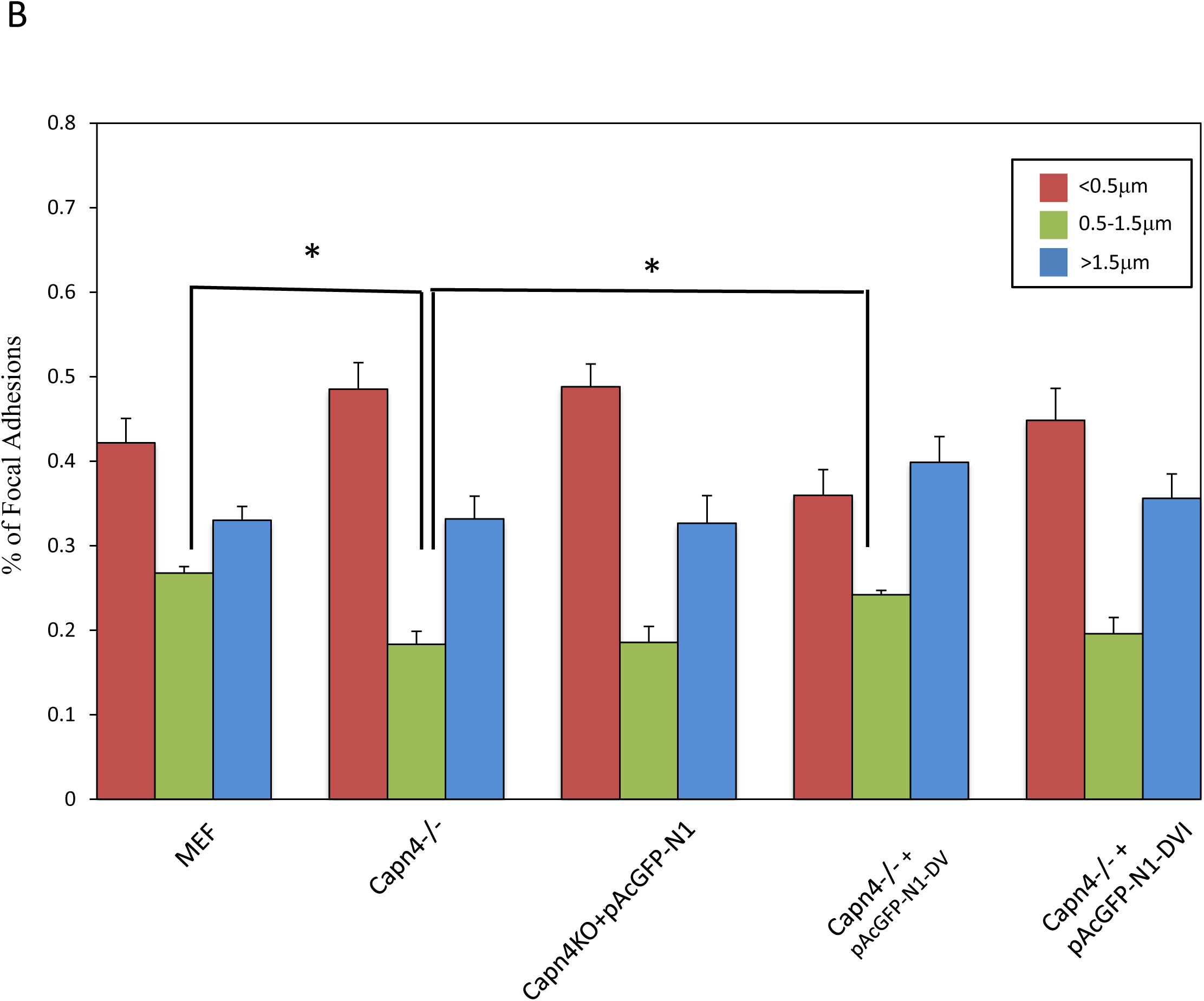
Overexpression of DV rescues the abnormal focal adhesion organization in *Capn4^−/−^* cells and promotes maturation of focal adhesions. *A*. Representative images show the immunofluorescence of focal adhesions with anti-vinculin antibody. Focal adhesions fail to mature into the cell body in *Capn4^−/−^* cells expressing DVI or an empty AcGFP-N1 plasmid compared to MEFs, while maturation of focal adhesions is rescued in *Capn4^−/−^* cells expressing DV (Mag. bar = 10μm). *B*. A Bar graph illustrates the percentage of average number of adhesions in terms of varying sizes in each of the cell lines. The numbers of focal adhesions were collected from 6 cells for each cell line. The number of nascent adhesions (0.5-1.5 sq.µm) is significantly reduced in *Capn4^−/−^* cells compared to wildtype MEFs, and overexpressing domain V increased this category significantly. Meanwhile, the overexpression of domain VI in *Capn4^−/−^* cells didn’t change this category significantly. The number of focal adhesions smaller than 0.5 sq.μm decreased significantly than *Capn4^−/−^* cells when domain V was expressed. When measuring focal adhesions larger than 1.5 sq.μm, no significant difference was observed between any two cell-lines. Statistical analysis was performed by student’s *t* test (* denotes p<0.05, ** denotes p<0.005, NS denotes a non-significant relationship).

### Overexpression of DV in Capn4^−/−^ Cells Promotes Cell Migration

Cell migration persistence and speed are affected by both biochemical and biophysical factors including dimension, stiffness, cell–cell and cell–matrix adhesion, traction forces, cytoskeletal polarity, and the capacity to degrade ECM by proteolytic enzymes, and so on. (Friedl and Wolf, 2010; Plotnikov et al., 2012; Wolf et al., 2013). It was previously observed that *Capn4^−/−^* cells have reduced migration rates (Dourdin et al., 2001; Undyala et al., 2008). To determine whether migration persistence and speed are affected by expressing either domain, *Capn4^−/−^* cells expressing either domain, *Capn4^−/−^* cells, and MEFs were seeded onto fibronectin coated glass coverslips and observed for 2 hours. Cell migration rates and persistence were calculated based on the locomotion of the nuclei. As expected, *Capn4^−/−^* cells migrated at a lower linear speed (0.50 μm/min) than MEFs (0.76 μm/min). Unlike *Capn4^−/−^* cells, *Capn4^−/−^* cells expressing DV migrated significantly faster (0.65 μm/min) than *Capn4^−/−^* cells in comparison to *Capn4^−/−^* cells expressing DVI (0.47 μm/min) (Fig. 4, *A*). In terms of persistence, no significance was observed between these two cell lines (Fig. 4, *B*). Together, these findings demonstrate that expressing DV in *Capn4^−/−^* cells rescues the migration speed defect, which is consistent with the observation that it also rescues the focal adhesion organization and traction force production.

**FIGURE 4.**
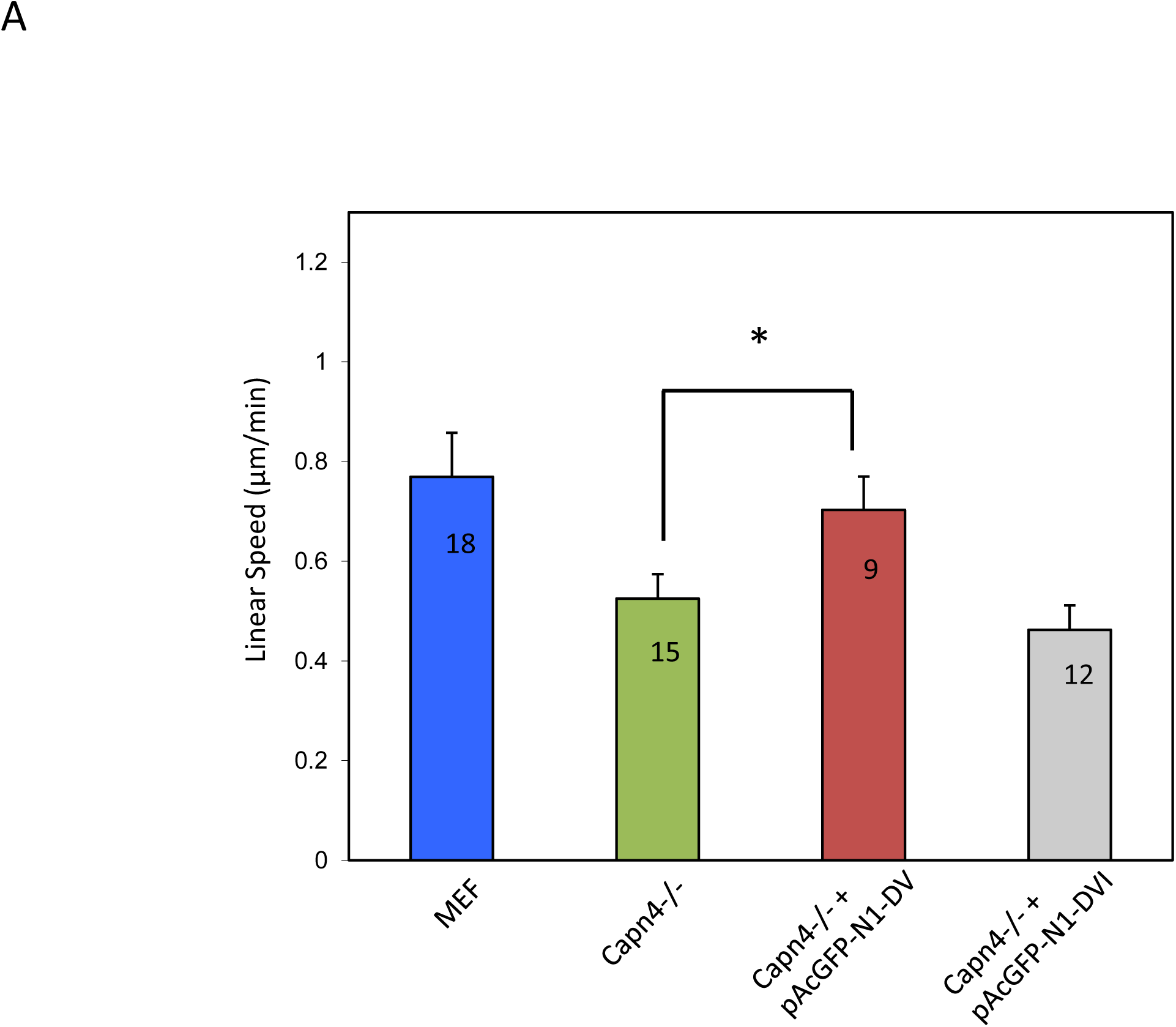
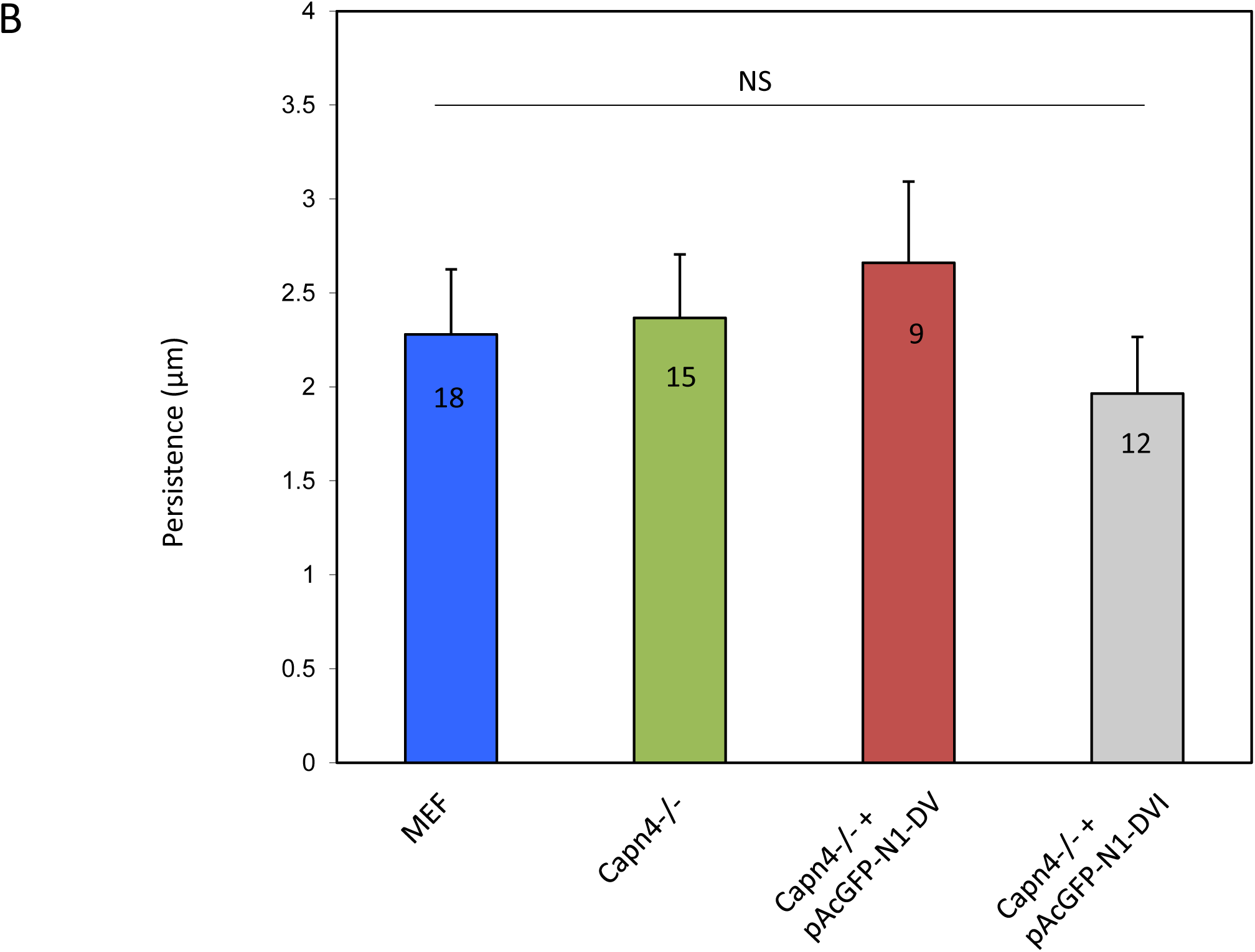
Overexpression of DV promotes cell migration speed in *Capn4^−/−^* cells but not the persistence. *A*. The bar graph represents the average of migration speed of MEFs, *Capn4^−/−^* cells, *Capn4^−/−^* cells expressing DV, and *Capn4^−/−^* cells expressing DVI. MEF cells migrated significantly faster than *Capn4^−/−^* cells. Expressing DV but not DVI in *Capn4^−/−^* cells increases the migration speed significantly compared to control *Capn4^−/−^* cells. *B*. The persistence of migration in each cell line was also calculated. No significant difference in persistence was observed between any two cell-lines. Numbers of cells used were marked in bar graphs. Statistical analysis was performed by student’s *t* test (* denotes p<0.05, ** denotes p<0.01, NS denotes a non-significant relationship).

### Overexpression of DVI Restores the Proteolytic Activity in Capn4^−/−^ Cells

It was previously reported that knocking-out of the calpain small subunit diminishes the proteolytic activity of the holoenzyme (Dourdin et al., 2001; Undyala et al., 2008). Since domain VI dimerizes with the calpain large subunit through its fifth EF-hand motif (Goll, 2003), we wanted to ask whether restoring DVI to *Capn4^−/−^* cells would restore the proteolytic activity of the holoenzyme. The calpain activity assay was performed using a calpain activity fluorometric assay kit with *Capn4^−/−^* cells expressing DV, DVI, full length *Capn4*, *Capn4^−/−^* cells, and MEFs. As shown in figure 5, *Capn4^−/−^* cells contain significantly lower levels of proteolytic activity in comparison with MEFs. However, in *Capn4^−/−^* cells expressing DVI and full length *Capn4*, this loss of proteolytic activity was restored. The same phenomena was not found in *Capn4^−/−^* cells expressing DV. This activity assay revealed that the presence of DVI is critical in the generation of holoenzyme proteolytic activity. The dimerization between the calpain small and large subunit might play a critical role in this process, which requires further experimentation to determine.

**FIGURE 5.**
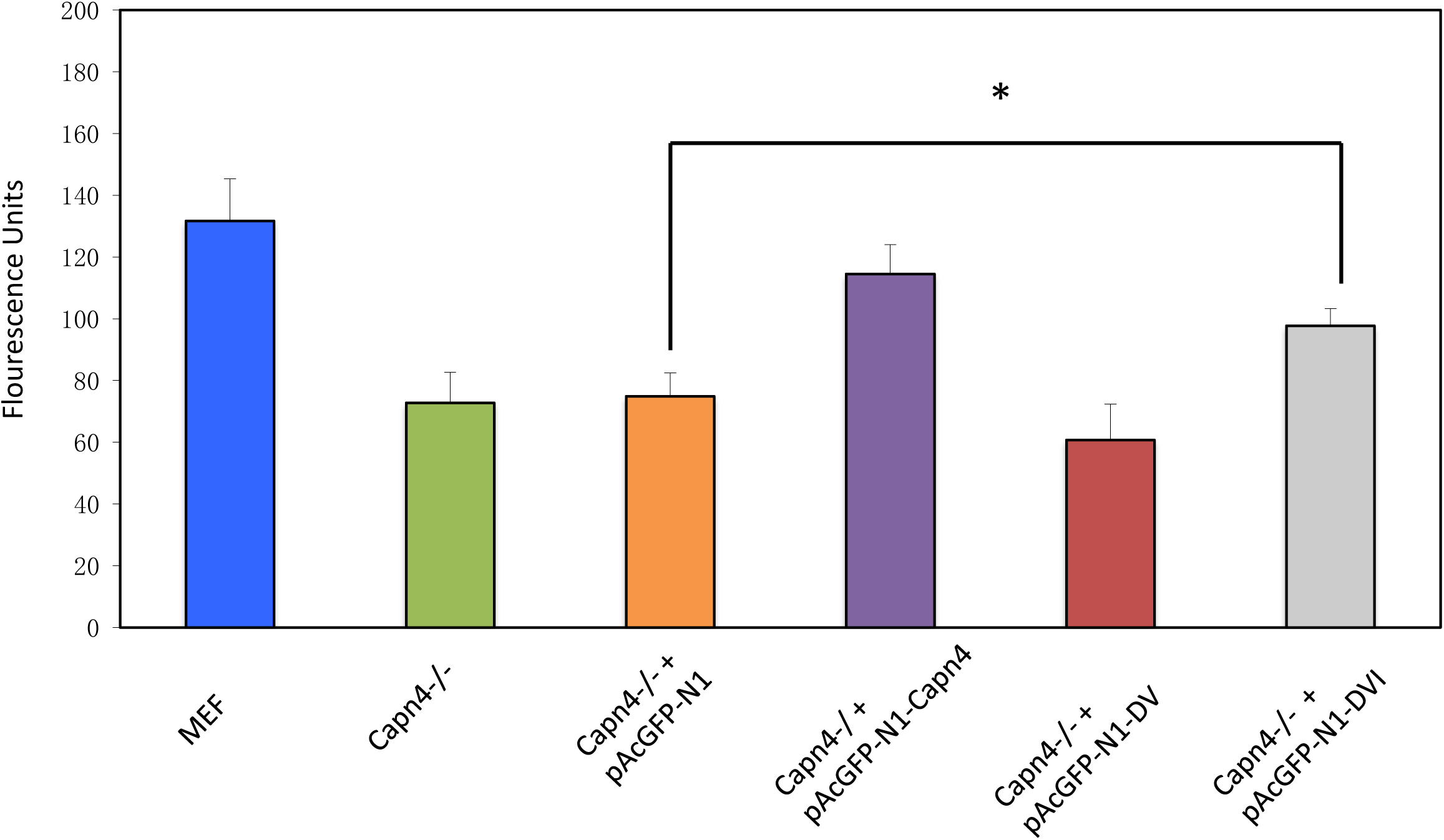
The bar graph indicates the relative fluorescence units representing calpain activity levels in each cell line obtained using the Biovision assay kit. The calpain activity was significantly reduced in *Capn4^−/−^* cells compared to MEFs. Expressing DVI in *Capn4^−/−^* cells significantly elevated the calpain activity compared to control *Capn4^−/−^* cells while expressing DV did not have the same effect. Statistical analysis was performed by student’s *t* test (* denotes p<0.05, ** denotes p<0.01, NS denotes a non-significant relationship).

## DISCUSSION

The proteolytic function of calpain holoenzymes play critical physiological roles, including cytoskeletal remodeling, cell differentiation, apoptosis, signal transduction, and so on (Carafoli and Molinari, 1998; Sato and Kawashima, 2001). Domain II on the large subunit contains the active site and is the only cysteine protease domain of the holoenzyme. For many years, the calpain small subunit’s function has been believed to be limited to assisting the proteolytic process of calpain holoenzymes (Goll et al., 2003). Previous research indicated that in *Capn4* deficient fibroblasts, the production of traction forces is impaired (Dourdin et al., 2001). One would expect *Capn4* deficient fibroblasts to generate consistent phenotypes similar to the ablation of *Capn1* and *Capn2* genes based on the canonical concept of the calpain small subunit’s function. However, our study attributes the reduction of traction forces in *Capn4^−/−^* cells solely to the calpain small subunit. We found that *Capn4* disruption inhibits both traction force production and mechanosensing, whereas inhibition of *Capn1* and/or *Capn2* impairs only mechanosensing but not traction force production (Undyala et al., 2008). These findings suggest a novel protease independent function for the calpain small subunit. To understand the mechanism that regulates traction force production through the calpain small subunit, we evaluated the magnitude of traction force and mechanosensing when each domain of the calpain small subunit was overexpressed in *Capn4^−/−^* fibroblasts. The most intriguing finding is that only the overexpression of domain V was sufficient to rescue the deficient traction force production in *Capn4^−/−^* cells, and that the overexpression of domain VI, but not domain V, restored the ability to sense the applied force onto the external substrates.

Domain V of the calpain small subunit is Gly rich with two stretches of 11 and 20 Gly residues and contains a common motif (GTAMRILGGVI) at the C-terminus. Numerous studies have suggested a phospholipid binding property for this common motif (Brandenburg et al., 2002; Daman et al., 2001), although the presence of this binding and attributed function are controversial (Goll et al., 2003). It has been suggested that the binding between domain V and phospholipids brings the holoenzyme close to the cell membrane in order to decrease the Ca^2+^ requirement for m-calpain activation (Johnson and Guttmann, 1997). One possibility is that this interaction is also important for domain V to position close to adhesion structures and initiate a protease independent pathway to regulate traction force production. The calpain holoenzyme undergoes a fast autolysis process during which 91 NH_2_-terminal amino acids are removed sequentially to produce 26-27kDa, then 22-23kDa, and finally, 18kDa autolytic fragments (Goll et al., 2003). Whether autolysis still occurs and if the rescue of the traction force production requires the presence of the whole domain V, or just the fragments released by autolysis, is unclear.

While overexpression of domain V rescues the traction force production in *Capn4^−/−^* cells, it also rescues the abnormal focal adhesion arrangement and maturation observed in these cells. More focal adhesions mature into the center of cells, and a higher percentage of focal adhesions fall into the category of nascent focal adhesions (0.5-1.5 sq.μm). This is consistent with previous observations that traction forces modulate lamellipodial extension, maturation of focal adhesions, and translocation of focal adhesions toward interior regions of the cell (Ridley et al., 2003), and that nascent adhesions generate stronger forces (Beningo et al., 2001). Multiple parameters are known to modulate the speed and persistence of cell migration, such as adhesiveness, strength of traction stress, the capacity to degrade ECM by proteolytic enzymes, and so on (Friedl and Wolf, 2010; Plotnikov et al., 2012; Wolf et al., 2013). In concert with elevated level of traction forces and rescue of focal adhesion arrangements, the cell migration speed was increased by overexpressing domain V in *Capn4^−/−^* cells on substrates in our study. Calpain 4 has been suggested to regulate the secretion of galactin-3 by indirectly mediating tyrosine phosphorylation (Menon et al., 2011). A possible mechanism for this is that calpain 4 mediates the secretion of galectin-3 indirectly through the binding of domain V with other interacting proteins or the cell membrane. Galectin-3 in the extracellular environment leads to clustering and activation of integrins (Goetz et al., 2008). Activated integrins then activate more downstream signaling proteins that ultimately lead to increased levels of traction forces, cell migration speed, and adhesion maturation.

Domain VI of the calpain small subunit (specifically, the COOH-terminus) is a calmodulin-like domain and contains five EF-hand motifs, the fifth of which heterodimerizes with the large subunits to form holoenzymes (Goll et al., 2003). It is already known that sensing the applied force requires functional calpain 1, 2, and 4 (Undyala et al., 2008). In concert, our results indicate that expressing domain VI restores the ability for *Capn4^−/−^* cells to sense the applied pushing force onto the substrate. Given the evidence that the large and small subunits remain associated when calpain is active (Johnson and Guttmann, 1997), this binding between the large and small subunit might play an important role in regulating mechanosensing. Moreover, since expression of domain VI also restores the calpain protease activity in *Capn4^−/−^* cells, as shown by our study, it is possible that mechanosensing is related to the holoenzyme’s protease function. Previous research identified an interaction between αPIX and calpain 4 (Rosenberger and Kutsche, 2005). αPIX interacts with the C-terminus of calpain 4 at the triple domain of SH3-DH-PH, and the integrity of the triple domain is necessary for efficient interaction between two proteins. This interaction is required for a cell to spread since the impairment of cell spreading resulting from inhibition of m-calpain in CHO-K1 cells can be rescued by overexpression of αPIX wild-type or GEF activity-deficient mutant, but not by the αPIX mutant in which domain DH is missing. These results also suggest that αPIX acts downstream of calpain to regulate cell spreading (Rosenberger et al., 2005). Based on these findings, αPIX is highly likely to be implicated in the mechanosensing pathway. Upon engagement to the ECM proteins, integrins are activated and cluster to form complexes. At the same time, structural and signaling molecules are recruited intracellularly to these early integrin clusters in which β1 integrin, ILK, calpain proteases, β-parvin, α-actinin, and αPIX are present but without paxillin and vinculin. These clusters might then allow mechanosensing to occur and may or may not require the GEF exchange activity of αPIX (Bialkowska et al., 2000; Rosenberger et al., 2005; Schoenwaelder and Burridge, 1999). These results are also consistent with our finding that mechanosensing in *Capn4^−/−^* cells could not be restored when only domain V of calpain 4 is overexpressed. There may be unidentified proteins containing the triple domain that interacts with calpain 4, either directly or indirectly, to mediate mechanosensing signal transduction.

In summary, we have found that the calpain small subunit not only plays a role in traction force production, which is beyond the regulatory function for the holoenzyme activity, but also that this function is only mediated through domain V. Meanwhile, it was also discovered that mechanosensing to localized forces is mediated through domain VI, but not domain V. This functional segregation is the first observation that both the traction force production and mechanosensing to localized mechanical forces are regulated through different domains of the same protein. This study provides new insight into the mechanism involving the calpain small subunit that regulates the generation of traction forces and the coordinate series of events that occur during cell migration.

## ACKNOWLEDGEMENTS

The authors wish to thank Drs. Gasparski, Menon and Jang for assistance with writing the manuscript. All experiments were performed by BQH. Portions of this work were developed from the dissertation of Dr. Bingqing Hao. Funding for this project was initially provided to KAB through NIH R01-GM084248 and start-up funds from Wayne State University.

---

**FIGURE S.1.**
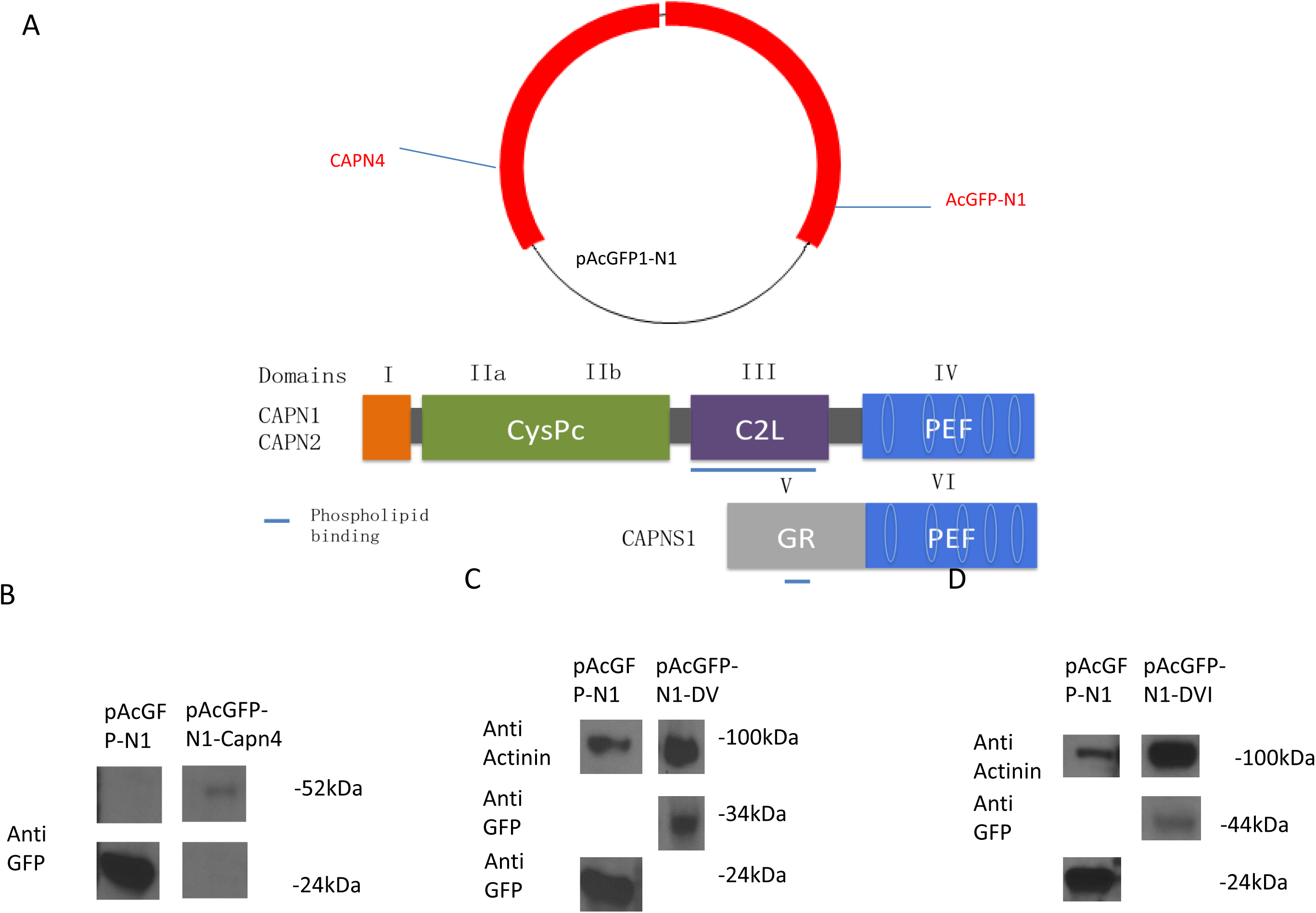
Over-expression of domain V or VI of *Capn4* in *Capn4^−/−^* cells. *A*, a schematic diagram illustrating the insertion of either DV or DVI of *Capn4*, or full-length *Capn4* into the plasmid pAcGFP1-N1 (Clontech). *B*, over-expression of DV, DVI, and full-length *Capn4* in *Capn4^−/−^* cells was confirmed by Western blots. Overexpression of DV is shown in the *left panel*, DVI in the *middle panel*, and full-length *Capn4* in the *right panel*.

## REFERENCES

1. Alexandrova AY, Arnold K, Schaub S, Vasiliev JM, Meister JJ, Bershadsky AD, Verkhovsky AB. (2008) Comparative dynamics of retrograde actin flow and focal adhesions: formation of nascent adhesions triggers transition from fast to slow flow. PLoS ONE 3(9). [PMID:18800171]

2. Beningo KA, Dembo M, Kaverina I, Small JV, Wang YL. (2001) Nascent focal adhesions are responsible for the generation of strong propulsive forces in migrating fibroblasts. J Cell Biol. 153(4): 881–8

3. Beningo, K.A. and Y.-L. Wang. (2002) Flexible substrata for the detection of traction forces. Trends in Cell Biol. 112(2), 79–84

4. Bialkowska K, Kulkarni S, Du X, Goll DE, Saido TC, Fox JE. (2000) Evidence that beta3 integrin-induced Rac activation involves the calpain-dependent formation of integrin clusters that are distinct from the focal complexes and focal adhesions that form as Rac and RhoA become active. J Cell Biol. 151(3):685–96

4. Bhatt, A., Kaverina, I., Otey, C. and Huttenlocher, A. (2002) Regulation of focal complex composition and disassembly by the calcium-dependent protease calpain. J. Cell Sci.115, 3415–3425

5. Brandenburg K, Harris F, Dennison S, Seydel U, Phoenix D. (2002) Domain V of m-calpain shows the potential to form an oblique-orientated alpha-helix, which may modulate the enzyme’s activity via interactions with anionic lipid. Eur J Biochem. 269(22): 5414–22

6. Carafoli E, Molinari M. (1998) Calpain: a protease in search of a function. Biochem Biophys Res Commun. 247(2): 193–203

7. Chang SS, Guo WH, Kim Y, Wang YL. (2013) Guidance of cell migration by substrate dimemsion. Biophys J. 104(2): 313–21

8. Chrzanowska-Wodnicka M, Burridge K. (1996) Rho-stimulated contractility drives the formation of stress fibers and focal adhesions. J Cell Biol.133(6): 1403–15

9. Crawford C, Brown NR, Willis AC. (1990) Investigation of the structural basis of the interaction of calpain II with phospholipid and with carbohydrate. Biochem J. 265(2): 575–9

10. Daman A, Harris F, Biswas S, Wallace J, Phoenix DA. (2001) A theoretical investigation into the lipid interaction of m-calpain. Mol Cell Biochem. 223(1-2): 159–63

11. Dembo, M. and Wang, Y.-L. (1999) Stresses at the cell-to-substrate interface during locomotion of fibroblasts. Biophys J. 76, 2307–2316

12. Dourdin, N., Bhatt, A. K., Dutt, P., Greer, P., Arthur, J. S., Elce, J. S. and Huttenlocher, A. (2001) Reduced cell migration and disruption of the actin cytoskeleton in calpain deficient-embryonic fibroblasts. J. Biol. Chem. 276, 48382–48388

13. Ekaterina Papusheva and Carl-Philipp Heisenberg. (2010) Spatial organization of adhesion: force-dependent regulation and function in tissue morphogenesis. EMBO J. 29(16): 2753–2768

14. Engler AJ, Sen S, Sweeney HL, Discher DE. (2006) Matrix elasticity directs stem cell lineage specification. Cell. 126(4): 677–89

15. Farshid Guilak, Daniel M. Cohen, Bradley T. Estes, Jeffrey M. Gimble, Wolfgang Liedtke, Christopher S. Chen. (2009) Control of stem cell fate by physical interactions with the extracellular matrix. Cell Stem Cell. Volume 5, Issue 1:17–26

16. Flevaris, P., Stojanovic, A., Gong, H., Chishti, A., Welch, E. and Du, X. (2007) A molecular switch that controls cell spreading and retraction. J. Cell Biol. 179, 553–565

17. Franco SJ, Huttenlocher A. (2005) Regulating cell migration: calpains make the cut. J Cell Sci. 118(Pt 17): 3829–38

18. Franco, S. J., Perrin, B. J. and Huttenlocher, A. (2004a). Isoform specific function of calpain 2 in regulating membrane protrusion. Exp. Cell Res. 299, 179–187

19. Franco, S. J., Rodgers, M. A., Perrin, B. J., Han, J., Bennin, D. A., Critchley, D. R. and Huttenlocher, A. (2004b) Calpain-mediated proteolysis of talin regulates adhesion dynamics. Nat. Cell Biol. 6, 977–983

20. Goetz, J.G., B. Joshi, P. Lajoie, S.S. Strugnell, T. Scudamore, L.D. Kojic, and I.R. Nabi. 2008. Concerted regulation of focal adhesion dynamics by galectin-3 and tyrosine-phosphorylated caveolin-1. J Cell Biol. 180:1261–1275.

20. Goll, D. E., Thompson, V. F., Li, H., Wei, W. and Cong, J. (2003) The calpain system. Physiol. Rev. 83, 731–801

21. Hynes RO. (2002) Integrins: bidirectional, allosteric signaling machines. Cell. 110(6): 673–87

22. Huttenlocher A, Palecek SP, Lu Q, Zhang W, Mellgren RL, Lauffenburger DA, Ginsberg MH, Horwitz AF. (1997) Regulation of cell migration by the calcium-dependent protease calpain. J Biol Chem. 272(52): 32719–22

23. Johnson GV, Guttmann RP. (1997) Calpains: intact and active? Bioessays. 19(11): 1011–8

24. Katarina Wolf, Mariska te Lindert, Marina Krause, Stephanie Alexander, Joost te Riet, Amanda L. Willis, Robert M. Hoffman, Carl G. Figdor, Stephen J. Weiss, Peter Friedl. (2013) Physical limits of cell migration: Control by ECM space and nuclear deformation and tuning by proteolysis and traction force. J Cell Biol. 201(7): 1069–1084

25. Lauffenburger DA, Horwitz AF. (1996) Cell migration: a physically integrated molecular process. Cell. 84(3): 359–69

26. Liedtke W, Kim C. (2005) Functionality of the TRPV subfamily of TRP ion channels: add mechano-TRP and osmo-TRP to the lexicon. Cell Mol Life Sci. 62(24): 2985–3001

27. Imajoh S, Kawasaki H, Suzuki K. (1986) The amino-terminal hydrophobic region of the small subunit of calcium-activated neutral protease (CANP) is essential for its activation by phosphatidylinositol. J Biochem. 99(4): 1281–4

28. Marganski, W.A., Dembo, M. and Wang, Y.-L. (2003) Measurements of cell-generated deformations on flexible substrata using correlation-based optical flow. Meth. Enzymol. 361, 197–121

29. Mamoune A, Luo JH, Lauffenburger DA, Wells A. (2003) Calpain-2 as a target for limiting prostate cancer invasion. Cancer Res. 63(15): 4632–40

30. Mathew E. Berginski, Eric A. Vitriol, Klaus M. Hahn, and Shawn M. Gomez. (2011) High-resolution quantification of focal adhesion spatiotemporal dynamics in living cells. PLoS One. 6(7): e22025

31. Menon, S., and K.A. Beningo. (2011) Cancer cell invasion is enhanced by applied mechanical stimulation. PLoS One. 6:e17277

32. Peter Friedl and Stephanie Alexande. (2011) Cancer invasion and the microenvironment: plasticity and reciprocity. Cell. 147(5): 992–1009

33. Peter Friedl, Katarina Wolf. (2010) Plasticity of cell migration: a multiscale tuning model. J Cell Biol. 188(1): 11–19

34. Pirta Hotulainen and Pekka Lappalainen. (2006) Stress fibers are generated by two distinct actin assembly mechanisms in motile cells. J Cell Biol. 173(3): 383–394

35. Potter DA, Tirnauer JS, Janssen R, Croall DE, Hughes CN, Fiacco KA, Mier JW, Maki M, Herman IM. (1998) Calpain regulates actin remodeling during cell spreading. J Cell Biol. 141(3): 647–62

36. Ridley AJ, Schwartz MA, Burridge K, Firtel RA, Ginsberg MH, Borisy G, Parsons JT, Horwitz AR. (2003) Cell migration: integrating signals from front to back. Science. 302 (5651): 1704–9

37. Rosenberger, G., A. Gal, and K. Kutsche. 2005. AlphaPIX associates with calpain 4, the small subunit of calpain, and has a dual role in integrin-mediated cell spreading. J Biol Chem. 280:6879–6889

37. Sato K, Kawashima S. (2001) Calpain function in the modulation of signal transduction molecules. Biol Chem. 382(5): 743–51.

38. Schoenwaelder, S.M., and K. Burridge. (1999) Bidirectional signaling between the cytoskeleton and integrins. Curr. Opin. Cell Biol. 11: 274–286

39. Sergey V. Plotnikov, Ana M. Pasapera, Benedikt Sabass, Clare M. Waterman. (2012) Force Fluctuations within Focal Adhesions Mediate ECM-Rigidity Sensing to Guide Directed Cell Migration. Cell. 151(7): 1513–27

40. V. Undyala, Micah Dembo, Katherine Cembrola, Benjamin J. Perrin, Anna Huttenlocher, John S. Elce, Peter A. Greer, Yu-li Wang, Karen A. Beningo. (2008) The calpain small subunit regulates cell-substrate mechanical interactions during fibroblast migration. J Cell Sci. 121(Pt 21): 3581–3588

